# Investigating real-life emotions in romantic couples: a mobile EEG study

**DOI:** 10.1101/2020.08.20.259796

**Authors:** Julian Packheiser, Gesa Berretz, Noemi Rook, Celine Bahr, Lynn Schockenhoff, Onur Güntürkün, Sebastian Ocklenburg

## Abstract

The neural basis of emotional processing has been largely investigated in constrained spatial environments such as stationary EEGs or fMRI scanners using highly artificial stimuli like standardized pictures depicting emotional scenes. Typically, such standardized experiments have low ecological validity and it remains unclear whether their results reflect neuronal processing in real-life affective situations at all. Critically, emotional situations do not only encompass the perception of emotions, but also behavioral components associated with them. In this study, we aimed to investigate real-life emotions by recording couples in their homes using mobile EEG technology during embracing, kissing and emotional speech. We focused on asymmetries in affective processing as emotions have been demonstrated to be strongly lateralized in the brain. We found higher alpha and beta power asymmetry during kissing and embracing on frontal electrodes during emotional kisses and speech compared to a neutral control condition indicative of stronger left-hemispheric activation. In contrast, we found lower alpha power asymmetry at parieto-occipital electrode sites in the emotional compared to the neutral condition indicative of stronger right-hemispheric activation. Our findings are in line with models of emotional lateralization that postulate a valence-specific processing over frontal cortices and right-hemispheric dominance in emotional processing in parieto-occipital regions. Overall, we could thus support theories of emotional asymmetries which suggest that affective processing is not uniformly lateralized across the brain using a highly ecologically valid paradigm.

## Introduction

One of the most intriguing questions in neuroscience revolves around how and where emotions are processed in the brain. More than 100 years ago, studies in patients with unilateral right-hemispheric lesions demonstrated that emotional processing seems to be lateralized in the brain as these patients were impaired in their ability to express their emotions (Mills, 1912). These results were complemented by a large number of behavioral results both from healthy and patient cohorts supporting the notion that emotions are asymmetrically processed in the brain (e.g., Borod et al., 1998; Landis, Assal, & Perret, 1979; Ley & Bryden, 1979; Suberi & McKeever, 1977). With the emergence of brain recording and neuroimaging techniques such as the EEG or fMRI (e.g., Davidson, Ekman, Saron, Senulis, & Friesen, 1990; Ekman, Davidson, & Friesen, 1990; Wager, Phan, Liberzon, & Taylor, 2003), more specific statements about where affective states are processed within cortical and subcortical regions could be made. For cortical emotional processing, especially frontal alpha power has been strongly associated with changes in affective state and emotional regulation (Allen, Keune, Schönenberg, & Nusslock, 2018; Reznik & Allen, 2018). For example, Hannesdóttir and colleagues (2010) investigated relative left frontal asymmetry (rLFA) and found that reduced rLFA in children was predictive of impaired emotional regulation and stronger physiological responses to emotional stimuli. Furthermore, rLFA serves as a suitable predictor for individual differences in emotional expression as well as regulation (Minnix & Kline, 2004; Papousek, Harald Freudenthaler, & Schulter, 2011).

Regarding the nature of emotional lateralization, two theories have been dominant in research on asymmetries of emotion processing. These theories are known as the right hemisphere hypothesis (RHH) and the valence model (VM) of emotional processing. The RHH postulates that all emotions regardless of valence are processed in the right hemisphere (Gainotti, 2019). In contrast, the VM claims that positive emotions are dominantly processed in the left hemisphere whereas negative emotions are processed in the right hemisphere (Prete, Laeng, & Tommasi, 2014). Both theories have received much support from behavioral, electrophysiological as well as neuroimaging studies (for review, see Demaree, Everhart, Youngstrom, & Harrison, 2005). The nature of emotional lateralization in the brain therefore remains rather inconclusive to this day. Despite the richness of the neuroscientific literature on emotions, the vast majority of published papers in this field share a common issue: It is largely unclear to what extent the used paradigms elicit neuronal processes that actually resemble the processes during real-life emotional encounters. A potential reason for this heterogeneity in the literature might be the lack of ecological validity in emotion research as the most prevalent method of positive or negative emotional induction is via movies, pictures or music (e.g. Gross & Levenson, 1995; Hausmann, Hodgetts, & Eerola, 2016; Hewig et al., 2005; Uhrig et al., 2016). Thus, emotions are largely only perceived during experimental paradigms. However, merely perceiving emotions might not be sufficient for a valid measurement of the underlying neural substrates. Real-life emotions comprise both the feeling itself as well as a preparation for and performance of an adequate action associated with the felt emotion (Güntürkün, 2019). This for example involves a behavioral expression such as avoidance behavior if someone experiences fear, or approach behavior if someone experiences happiness (for review, see Ocklenburg, Berretz, Packheiser, & Friedrich, 2020). The lack of a behavioral component in laboratory settings challenges the external validity of experimental results and makes it difficult to transfer findings to, for example, mood disorders for which the associated behavior is of paramount importance.

In conventional settings, this behavioral component is unfortunately difficult to realize either due to the experimental design or the environment in which it takes place, for example in an fMRI scanner or during EEG recordings. The past decade however brought forth novel techniques such as mobile EEGs (Gramann et al., 2011; Vos & Debener, 2014), mobile fNIRS (Holtzer et al., 2011; Quaresima & Ferrari, 2019) and less constraining MEGs (Boto et al., 2018; Boto et al., 2019; Roberts et al., 2019). These technological advancements have been developed and optimized to allow for the investigation of neural correlates in settings of high ecological validity as the participants can move freely and be tested outside the lab. While the number of publications using these techniques are still sparse, a growing body of research is being generated converging neuroscientific research with real-life activities such as cycling (Scanlon, Townsend, Cormier, Kuziek, & Mathewson, 2019), walking (Lin et al., 2020), navigating over obstacles (Nordin, Hairston, & Ferris, 2019), or skateboarding (Robles et al., 2020). Recently, Packheiser and colleagues (2020) investigated the neural basis of hand and foot use while the participants were wearing a mobile EEG. They found that both alpha and beta frequency asymmetries were predictive of the participants’ handedness and footedness and that the neural signals could distinguish between limb preferences of individuals. Importantly, they found that the neural signals were unaffected by movement parameters during activities such as jumping or throwing and kicking balls. Thus, mobile EEGs provide a valuable tool to investigate the neural basis of human emotions and their associated behaviors in more natural settings.

A very prominent human behaviour that is usually executed in emotional settings is social touch. To convey our social intentions or emotional states to other humans, we strongly rely on the use of a variety of tactile interactions, especially in very intimate social relationships (Dunbar, 2010; Frith & Frith, 2007). Touch is the earliest sensory modality to fully develop during the lifespan (Maurer & Maurer, 1988) and is experienced from birth onwards by being cradled in the mother’s arms (Forsell & Åström, 2012). For that reason, social touch has been strongly associated with human development shaping attachment, emotional regulation and cognitive maturation (Cascio, Moore, & McGlone, 2019). Affective social touch has been demonstrated to be highly beneficial for the well-being and physical as well as mental health in humans as it reduces stress, blood pressure and can even protect from viral infections and allergic responses (Cohen, Janicki-Deverts, Turner, & Doyle, 2015; Floyd et al., 2009; Kimata, 2006; Light, Grewen, & Amico, 2005). Studies on the neural basis of social touch have indicated that especially limbic and orbitofrontal brain regions are activated when an experimenter or the romantic partner applies non-sexual pleasant tactile stimulation (Li et al., 2019; Morrison, 2016; Nummenmaa et al., 2016). Thus, there seems to be a strong overlap in cerebral processing of social touch and emotions indicating that social touch carries a strong affective component. However, as for studies investigating emotions, the experimental designs investigating the neural basis of social touch lack ecological validity as the application of gentle touch to e.g. the legs while lying perfectly still rarely occurs in real-life settings.

The aim of the present study was to investigate emotional lateralization in a setting with high ecological validity. To this end, we tested romantic partners in their home during both embracing and kissing while the participants were recorded using a mobile EEG system. We also investigated the neural correlates of emotional speech as this type of social interaction is fundamental to maintain a healthy and long-lasting relationship (Litzinger & Gordon, 2005). We focused on differences in asymmetrical processing in the alpha frequency band due to the pronounced role of frontal alpha asymmetries in emotional processing. Since beta power asymmetries have been demonstrated to be highly comparable in function to alpha power asymmetries in studies investigating motor preferences (Packheiser, Schmitz, Pan et al., 2020) and resting state oscillations (Ocklenburg et al., 2019), we also included beta power asymmetries as a dependent variable in our study.

## Methods

### Participants

A total of 32 individuals (16 females) took part in this study. The sample size was determined based on prior studies investigating frontal alpha asymmetries using within-subject designs producing reliable and moderate to large effects (e.g., Flo et al., 2011; Mikutta, Altorfer, Strik, & Koenig, 2012). There was no restriction regarding the sexuality of the participants, but all couples were heterosexual in the present study. Age of the participants ranged between 19 and 63 years (mean age = 29 years, SD = 14 years). Participants with neurological or psychiatric disorders were excluded from the study. The study was conducted in accordance with the declaration of Helsinki and was approved by a local ethics committee of the psychological faculty at Ruhr University Bochum. All participants gave written informed consent.

### Experimental task

The experimental paradigm consisted of conducting three behavioral tasks, i.e. embracing, kissing and listening to speech in both an emotional and neutral condition. The neutral condition served as a control condition and was conducted using identical movements to control for motor effects in the experiment. Testing took place in the participants’ homes to provide as much natural setting and ecological validity as possible. One participant was set up with the mobile EEG system while the other participant filled out demographic questionnaires and prepared an emotional text about fond memories and experiences with their partner. Importantly, the partner, that was not recorded, was also equipped with an electrode cap to reduce the awkwardness of only one partner wearing the cap. Next, the participants were instructed about the behavioral tasks and how to perform them appropriately by the experimenter and an illustrating photograph. After instructing the participants, the experimenter left the room to allow for privacy for the entire experimental procedure. The behavioral tasks looked as follows:
 
1. During the embracing condition, the participants were either asked to embrace each other from the front in the emotional condition (figure 1A), or embrace a body pillow in the neutral condition with the partner being absent (figure 1B). The participants were further instructed to avoid touching the electrode cap and move as little as possible during the embrace.
2. During the kissing condition, the participants were either asked to kiss each other on the lips in the emotional condition (figure 1C), or kiss their own hand during the neutral condition by forming a lip-like structure with the thumb and index finger with the partner being absent (figure 1D). Importantly, the participants were instructed not to use their tongue during the kiss in either condition to avoid strong motor artefacts.
3. During the speech condition, the participant wearing the mobile EEG system was either listening to the partner reading the previously prepared emotional text in the emotional condition (figure 1E), or was listening to a weather report that was recorded prior to the experimental session in the neutral condition with the partner being absent (figure 1F).

**Figure 1.**
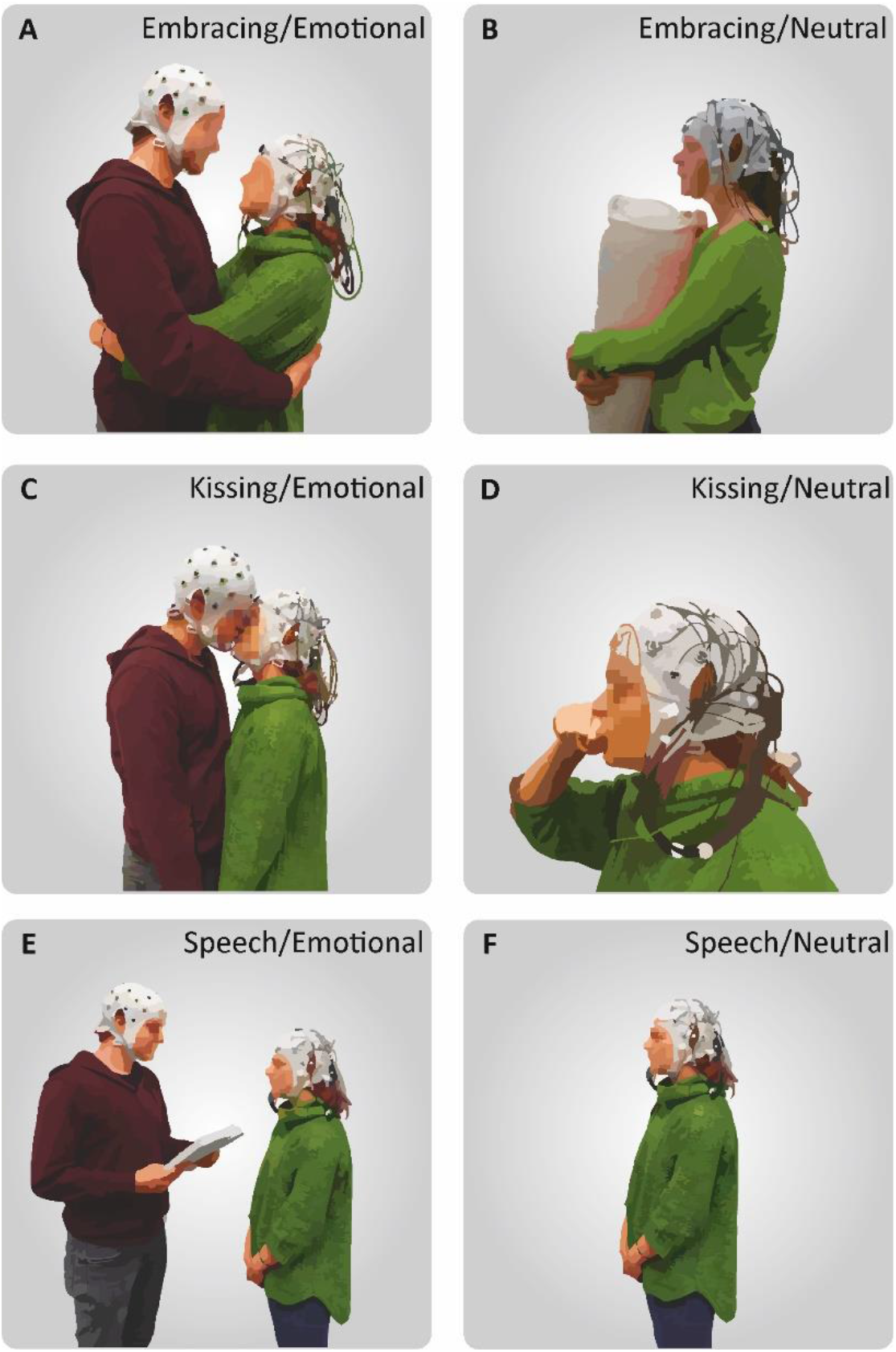
Depiction of the behavioral tasks. (A) During the emotional condition, the participant wearing the mobile EEG system embraced his/her partner in a frontal embrace. (B) During the neutral condition, the participant embraced a pillow with the partner absent. (C) In the emotional kissing condition, the partners kissed each other without using their tongue. (D) During the neutral condition, the recorded participant performed a kiss with his or her own hand. (E) In the emotional speech condition, the recorded participant listened to an emotional text written by their partner prior to the session about shared life experiences. (F) In the neutral speech condition, the recorded participant listened to a pre-recorded weather report without the partner.

The experimental design was counterbalanced so that the experiment could start with the neutral or emotional condition and with the male or female partner being recorded initially. Furthermore, during the emotional and neutral condition, the order of behavioral tasks was fully randomized. Each behavioral task was performed for one minute in total and the tasks were separated by an intertrial interval (ITI) of 30 second in accordance with the procedures used in Packheiser et al. (2020). During the ITI, the participant(s) received an additional pre-recorded auditory and visual instruction about the upcoming behavioral task. This was necessary as it was unknown to the participant(s) if the procedure started with embracing, kissing or speech due to the randomization procedure. The instructions were presented using the Presentation software (Neurobehavioral Systems Inc., CA, USA). There was a five minute break between the emotional and neutral condition to fill out the Positive and Negative Affect Schedule (PANAS), a questionnaire evaluating their positive and negative affective states by indicating the current emotional state on 10 positive and 10 negative items on a scale from 1 (Very slightly or not at all) to 5 (Extremely) (Crawford & Henry, 2004). The break was also used to allow for the partner to leave or to join (depending whether the recorded partner was in the emotional or neutral condition, respectively).

After one participant had completed the experimental procedure (three emotional and three neutral tasks), the recorded partner again filled out the PANAS so that the affective state was measured after both the emotional and neutral condition. Afterward, the roles of the partners switched, and the non-recorded partner went through the identical experimental protocol.

### EEG recording, preprocessing and analysis

EEG signals were obtained with a mobile EEG recording system (LiveAmp 32, Brain Products GmbH, Gilching, Germany). The LiveAmp 32 comprises 32 Ag-AgCL electrodes arranged in the international 10-20 system (C3/C4, FP1/FP2, Fz, F3/F4, F7/F8, FCz, FC1/FC2, FC5/FC6, FT9/FT10, T7/T8, CP1/CP2, CP5/CP6, TP9/TP10, Pz, P3/P4, P7/P8, Oz and O1/O2). The FCz electrode served as reference signal during data recording. All signals were amplified using a wireless amplifier (analog-to-digital conversion: 24-bit) and recorded using the Brain Vision analyzer software at a sampling rate of 1 kHz. Impedances were kept below 10 kHz during the recording session to ensure good signal quality. The EEG system furthermore comprised three acceleration sensors in the X (mediolateral axis), Y (anteroposterior axis) and Z (dorsoventral axis) direction located at the rear of the skull that recorded movements of the participants’ head.

Following data acquisition, the EEG signals were preprocessed offline in Brain Vision Analyzer (Brain Products GmbH, Gilching, Germany). The raw data files were band-pass filtered from 0.1 Hz (high pass) to 30 Hz (low pass) at 24 dB (octave). All signals were manually inspected for technical artifacts and channels of poor recording quality. Systematic artifacts, i.e. horizontal or vertical eye movements as well as pulse-associated signals, were removed via the application of an infomax independent component analysis (ICA). The reference channel (FCz) and channels of insufficient signal quality were recalculated via topographic interpolation.

After preprocessing, the individual tasks were first segmented across the entire trial duration (60 s) and then baseline corrected. The 500 ms prior to task onset were used as baseline signal. The large trial segment was then further divided into 58 non-overlapping segments of 1024ms duration each. Individual segments were excluded via an automatic artifact rejection if any of the following criteria were met: (1) voltage steps of 50 μV / ms, (2) amplitude differences of more than 200 μV within a 200 ms interval and (3) signal strength below 0.5 μV within a 100 ms interval. In a next step, we applied a current source density (CSD; (Peters & Servos, 1989) transformation to remove the reference potential from the filtered and segmented data. Finally, we used a Fast-Fourier transformation to decompose the oscillatory data into different its frequency bands (Hammond window of 10 %). Alpha oscillations were defined in the 8 – 13 Hz range. Beta frequencies were defined in the 13 – 30 Hz range. We then calculated the average power density (power per unit bandwidth) per electrode with a bilateral arrangement (C3/C4, FP1/FP2, F3/F4, F7/F8, FC1/FC2, FC5/FC6, FT9/FT10, T7/T8, CP1/CP2, CP5/CP6, TP9/TP10, P3/P4, P7/P8 and O1/O2) and extracted it for the three tasks across both conditions individually. In a final step, asymmetry indices (AIs) were computed between the electrode pairs using the following formula in accordance with Ocklenburg et al. (2019):

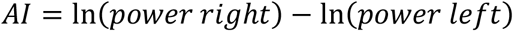

### Statistical analysis

Statistical analyses were conducted using SPSS (version 21, Chicago, Ilinois, USA). The PANAS scores were evaluated using a two-factorial repeated measures ANOVA with the factor valence (two levels: average score for all positive and all negative items) and the factor condition (two factors: emotional and neutral). Post hoc testing was performed using a Bonferroni correction. Neural data was analyzed separately for the three behavioral tasks. We investigated differences in AIs in the alpha and beta frequency band on all electrode-pairs for which they could be computed, i.e. for all non-central electrodes. We computed a two-factorial repeated measures ANOVA with each individual electrode pair as the first factor (14 levels: C3/C4, FP1/FP2, F3/F4, F7/F8, FC1/FC2, FC5/FC6, FT9/FT10, T7/T8, CP1/CP2, CP5/CP6, TP9/TP10, P3/P4, P7/P8 and O1/O2) and the experimental condition as second factor (two levels: emotional and neutral). Again, post hoc comparisons were conducted using a Bonferroni correction. To identify movement-related differences between the conditions, we extracted the acceleration sensor signals on the X-,Y- and Z-axis during the emotional and neutral condition. A two-factorial repeated measures ANOVA was performed with the factor orientation (three levels, X,Y and Z) and condition (two levels: emotional and neutral). Five participants were excluded from the PANAS analysis and one participant was excluded from the EEG analysis due to technical issues.

## Results

### Emotional induction

First, we investigated whether our emotional condition elicited more positive affective states compared to the neutral conditions using the PANAS scores. The results of the PANAS questionnaire were evaluated by comparing the average value of all positive and all negative items between the emotional and neutral condition in a 2×2 ANOVA. We found significant main effects of valence (F_(1,26)_ = 329.72, *p* < 0.001, *η*_*p*_^*2*^ = 0.93) and condition (F_(1,26)_ = 72.42, *p* < 0.001, *η*_*p*_^*2*^ = 0.74) with the positive items being rated higher (mean score = 2.75) than the negative items (mean score = 1.24) and the emotional condition receiving higher ratings (mean score = 2.35) compared to the neutral condition (mean score = 1.64). We found a significant interaction between item valence and the experimental conditions (F_(1,26)_ = 54.07, *p* < 0.001, *η*_*p*_^*2*^ = 0.68, see figure 2). Bonferroni-corrected post-hoc tests revealed significantly higher positive affect in the emotional (mean score = 3.46, SEM = 0.14) compared to the neutral condition (mean score = 2.05, SEM = 0.13, p < 0.001). Negative affect did not differ between the conditions (p > 0.250) and was basically absent in both the emotional (mean score = 1.23, SEM = 0.05) and the neutral condition (mean score = 1.25, SEM = 0.07).

**Figure 2.**
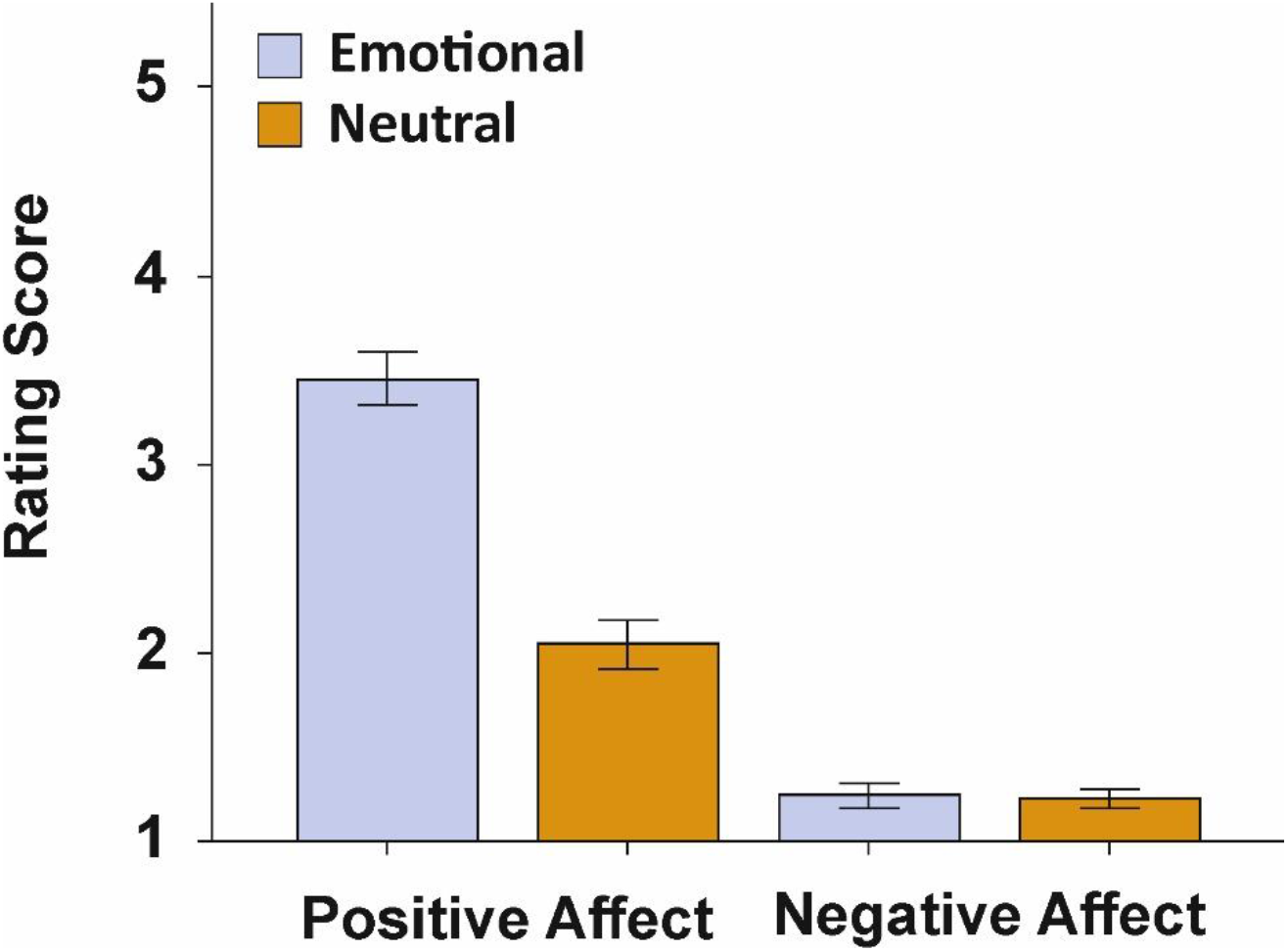
PANAS ratings indicating the momentary emotional state following the emotional or the neutral experimental condition. Error bars represent SEM.

### Alpha power asymmetries

To investigate differences in neural processing between the emotional and neutral condition, we investigated changes in AIs between the conditions for all three behavioral tasks individually in a 2 (factor condition) × 14 (factor electrode pair) ANOVA. For the embracing condition, we found neither a significant main effect of condition (F_(1,30)_ = 2.39, *p* = 0.133, *η*_*p*_^*2*^ = 0.07), nor a significant interaction between condition and electrode pairs (F_(13,390)_ = 0.61, *p* > 0.250, *η*_*p*_^*2*^ = 0.02, figure 3A). For the kissing condition, we found no significant main effect of condition (F_(1,30)_ = 0.18, *p* > 0.250, *η*_*p*_^*2*^ = 0.01), but a significant interaction between condition and electrode pairs (F_(13,390)_ = 1.95, *p* = 0.024, *η*_*p*_^*2*^ = 0.06). Bonferroni corrected post hoc testing revealed a significantly higher asymmetry index on the FP1/FP2 electrode pair in the emotional (mean μV^2^/Hz = 0.24, SEM = 0.10) compared to the neutral condition (mean μV^2^/Hz = −0.05, SEM = 0.10, *p* = 0.039, figure 3B). For the speech condition, we found no significant main effect of condition (F_(1,30)_ = 0.94, *p* > 0.250, *η*_*p*_^*2*^ = 0.03), but a significant interaction between condition and electrode pair (F_(13,390)_ = 2.28, *p* = 0.007, *η*_*p*_^*2*^ = 0.07). Bonferroni corrected post hoc testing revealed a significantly lower asymmetry index on the P7/P8 electrode pair in the emotional (mean μV^2^/Hz = −0.06, SEM = 0.08) compared to the neutral condition (mean μV^2^/Hz = 0.18, SEM = 0.06, *p* = 0.022, figure 3C).

**Figure 3.**
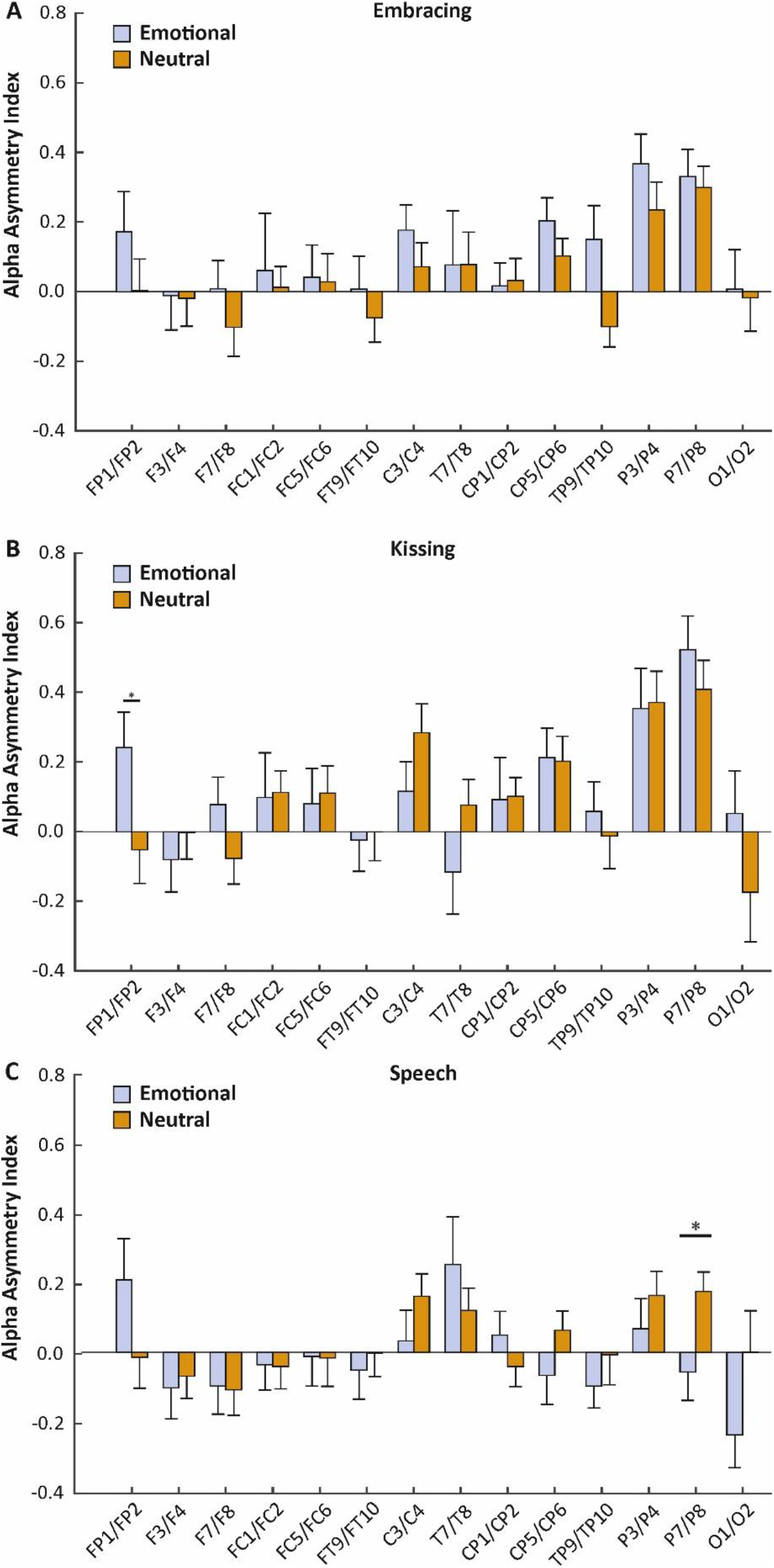
Electrode-specific analysis of alpha power asymmetries during the three behavioral tasks (A: Embracing, B: Kissing, C: Speech) in the emotional and neutral conditions. Error bars represent SEM.

### Beta power asymmetries

We repeated the analysis conducted for the alpha frequency band in the beta frequency band. For the embracing condition, we found neither a significant main effect of condition (F_(1,30)_ = 3.58, *p* = 0.068, *η*_*p*_^*2*^ = 0.11), nor a significant interaction between condition and electrode pairs (F_(13,390)_ = 1.49, *p* > 0.250, *η*_*p*_^*2*^ = 0.05, figure 4A). For the kissing condition, we found no significant main effect of condition (F_(1,30)_ = 1.27, *p* > 0.250, *η*_*p*_^*2*^ = 0.04), but a significant interaction between condition and electrode pairs (F_(13,390)_ = 2.67, *p* = 0.002, *η*_*p*_^*2*^ = 0.08). Bonferroni corrected post hoc testing revealed a significantly higher asymmetry index on the FP1/FP2 electrode pair in the emotional (mean μV^2^/Hz = 0.26, SEM = 0.09) compared to the neutral condition (mean μV^2^/Hz = −0.10, SEM = 0.09, *p* = 0.006). Furthermore, there was a significantly higher asymmetry index on the F7/F8 electrode pair in the emotional (mean μV^2^/Hz = −0.03, SEM = 0.10) compared to the neutral condition (mean μV^2^/Hz = −0.43, SEM = 0.14, *p* = 0.023, figure 4B). For the speech condition, we found no significant main effect of condition (F_(1,30)_ = 0.92, *p* > 0.250, *η*_*p*_^*2*^ = 0.03), but a significant interaction between condition and electrode pair (F_(13,390)_ = 2.41, *p* = 0.004, *η*_*p*_^*2*^ = 0.07). Bonferroni corrected post hoc testing revealed a significantly higher asymmetry index on the FP1/FP2 electrode pair in the emotional (mean μV^2^/Hz = 0.22, SEM = 0.10) compared to the neutral condition (mean μV^2^/Hz = −0.08, SEM = 0.11, *p* = 0.016). Furthermore, there was a significantly higher asymmetry index on the F7/F8 electrode pair in the emotional (mean μV^2^/Hz = −0.06, SEM = 0.13) compared to the neutral condition (mean μV^2^/Hz = −0.34, SEM = 0.12, *p* = 0.047, figure 4C).

**Figure 4.**
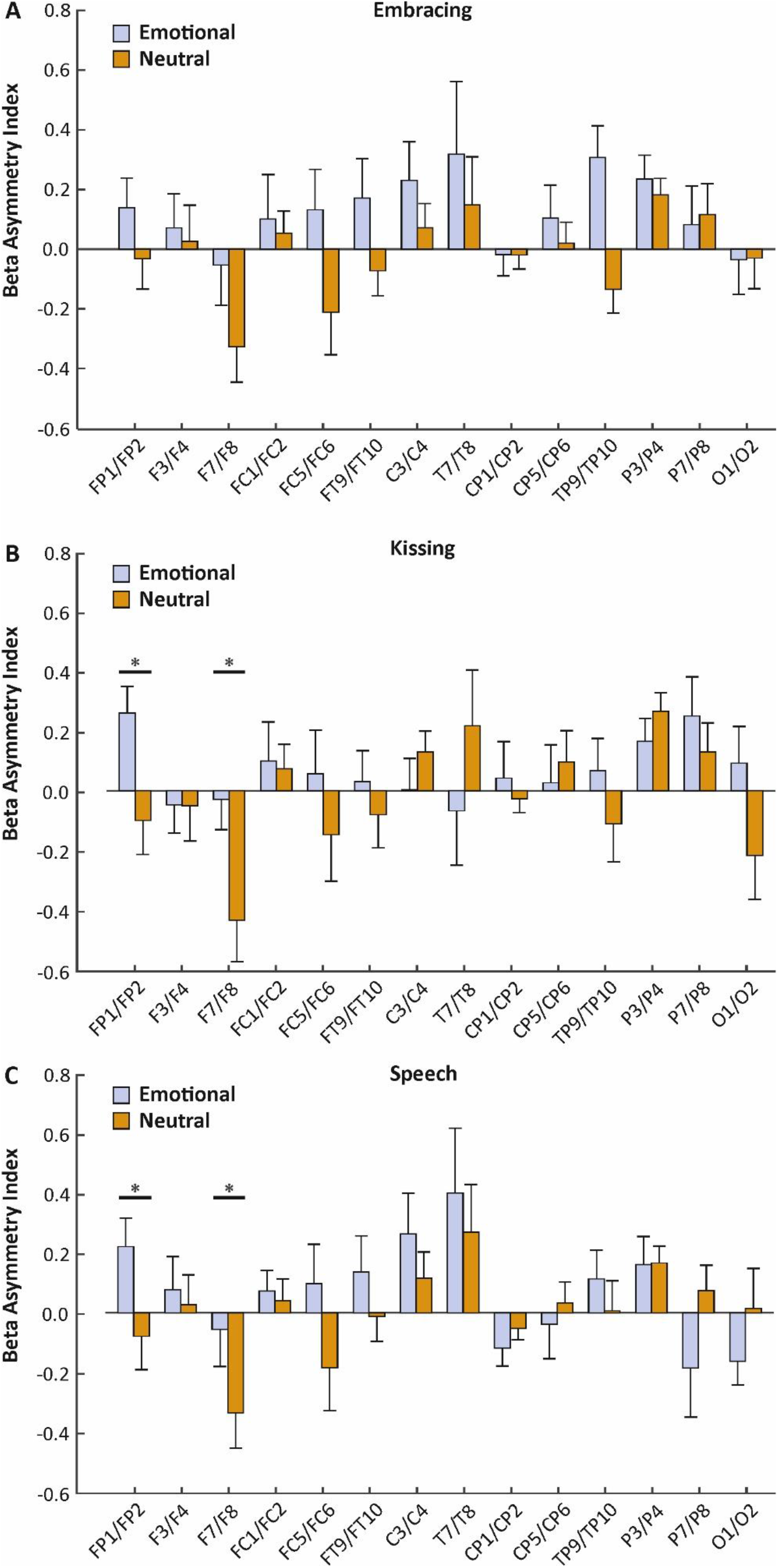
Electrode-specific analysis of beta power asymmetries during the three behavioral tasks (A: Embracing, B: Kissing, C: Speech) in the emotional and neutral conditions. Error bars represent SEM.

### Acceleration sensors

To identify whether the emotional condition was associated with stronger movement, we compared the acceleration sensor signals between the emotional and neutral condition for each behavioral task. To this end, we computed 2×3 ANOVA with the factors condition (two levels: emotional and neutral) and orientation (three levels: X, Y, and Z-axis). For embracing, we found no main effect of condition (F_(1,30)_ = 4.15, *p* = 0.051, *η*_*p*_^*2*^ = 0.12), nor an interaction with the movement orientation (F_(2,60)_ = 1.93, *p* = 0.154, *η*_*p*_^*2*^ = 0.06, see figure 5A). For kissing, the results were comparable as we also did not detect a main effect of condition (F_(1,30)_ = 1.01, *p* > 0.250, *η*_*p*_^*2*^ = 0.03), nor an interaction with the movement orientation (F_(2,60)_ = 0.08, *p* > 0.250, *η*_*p*_^*2*^ = 0.003, see figure 5B). Finally, the speech condition also did not yield any significant main effect of condition (F_(1,30)_ = 0.57, *p* > 0.250, *η*_*p*_^*2*^ = 0.02), nor an interaction with the movement orientation (F_(2,60)_ = 1.20, *p* > 0.250, *η*_*p*_^*2*^ = 0.04, see figure 5C). Grand averages of the movement signals split by frequency band (alpha, beta, gamma, delta) for the three behavioral tasks are depicted in SI figure 1.

**Figure 5.**
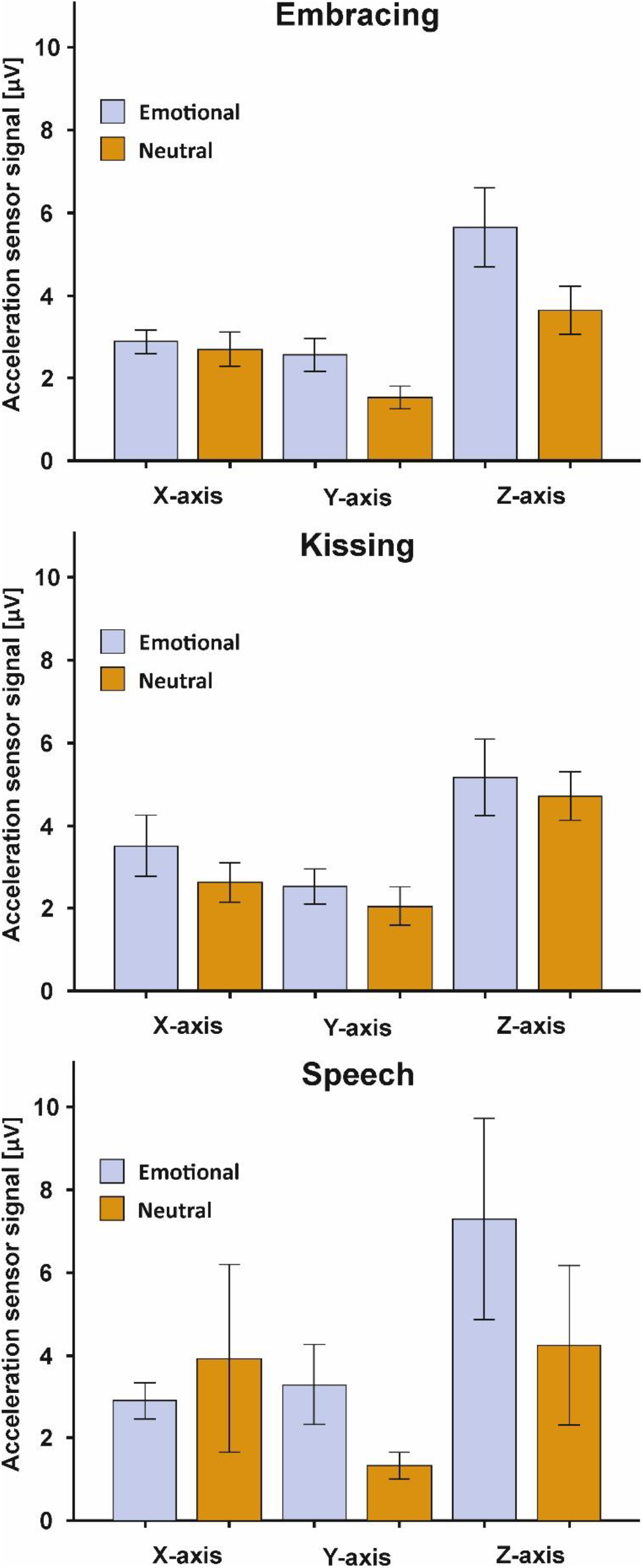
Acceleration sensor signals in the behavioral tasks for the X-.Y- and Z-axis orientation. Error bars represent SEM.

## Discussion

In the present study, we used a mobile EEG to record brain activity of romantic partners during affective social touch and emotional speech in their everyday environment to provide high ecological validity. We specifically focused on asymmetries in our analysis due to the pronounced lateralization of emotional processing in the brain. We found that the participants were in a more positive mood after they executed the behavioral tasks with their respective partner. On the neural level, we found a higher alpha AIs on the FP1/FP2 electrode pair in the emotional compared to the neutral condition during kissing. For speech, we found a lower alpha AI in the emotional compared to the neutral condition on the P7/P8 electrode pair. In the beta frequency band, we found higher AIs in the emotional compared to the neutral condition on the FP1/FP2 and F7/F8 electrode pair during both kissing and emotional speech. Movement signals did not differ between the emotional and neutral condition.

Increases in oscillatory alpha power have been strongly associated with functional inhibition, for example during visuospatial attention (Kelly, Gomez-Ramirez, & Foxe, 2009; Worden, Foxe, Wang, & Simpson, 2000), face recognition (Haegens, Osipova, Oostenveld, & Jensen, 2010) and working memory tasks (Jensen & Mazaheri, 2010). Alpha oscillations are hypothesized to be generated by rhythmic burst of local inhibitory GABAergic interneurons (Jensen & Mazaheri, 2010). Increases in AIs are therefore indicative of stronger right-hemispheric inhibition or increased left frontal activity whereas decreases in AIs reflect stronger left hemispheric inhibition or right-hemispheric activation. Thus, the frontal increase in alpha power asymmetries during emotional compared to neutral kisses indicates that frontal regions of the left hemisphere were more strongly activated in the presence of strong positive affect. These results are in line with the VM of emotional lateralization which postulates that positive emotions are processed in the left hemisphere and oppose predictions made by the RHH claiming that all emotions are processed in the right hemisphere irrespective of valence. Interestingly, previous behavioral research on the effects of emotional context on the lateralization of social behaviour has indicated that the RHH provides the overall best prediction to explain changes in laterality in emotional compared to neutral situations (Ocklenburg et al., 2018; Packheiser et al., 2019; Packheiser, Schmitz, Metzen et al., 2020). Prete and colleagues (2018) similarly found that behavioral and neural findings regarding hemispheric asymmetries were incongruent and do not necessarily correspond. It should be noted however that the VM and RHH are not necessarily mutually exclusive (Prete, Laeng, Fabri, Foschi, & Tommasi, 2015). Killgore and Yurgelun-Todd (2007) have proposed an integrative model postulating that the VM provides accurate predictions for anterior or frontal asymmetries whereas posterior or parietal asymmetries are more in line with predictions of the RHH. Since we could find lower asymmetry scores during emotional compared to neutral speech on the P7/P8 electrode pair indicating stronger right-hemispheric activation in the emotional condition, our results corroborate that emotional lateralization does not seem to be uniform across cortical brain regions but is rather region-specific.

For asymmetries in the beta frequency band, we found mostly comparable and sometimes even larger effects compared to the alpha band. Opposed to oscillatory alpha power, the functional role of the beta frequency band has been rather inconclusive. While some studies have suggested that beta activity is indicative of cognitive activation (Buschman, Denovellis, Diogo, Bullock, & Miller, 2012; Kamiński, Brzezicka, Gola, & Wróbel, 2012), beta power has also been suggested to be associated with the function of inhibitory interneuron networks indicating that alpha and beta activity share similar characteristics (Porjesz et al., 2002). In a previous mobile EEG study investigating alpha and beta asymmetries during motor execution, we also found that alpha and beta asymmetries were functionally similar and associated with inhibition (Packheiser, Schmitz, Pan et al., 2020). Ocklenburg et al. (2019) even found that alpha, beta, delta and theta oscillations were all significantly correlated indicating that there might be some common function underlying rhythmic brain activity in general. Our results provide further evidence that alpha and beta oscillations play a similar functional role that is likely linked to inhibition.

Opposed to the kissing and speech condition, we could not find any effects in the alpha or beta frequency band for the embracing condition. Given that the effects in the embracing condition were very small, it seems unlikely that the lack of significant results in this condition were due to a lack of power. However, there might have been other contributing factors that could explain this difference between the behavioral tasks. One potential explanation relates to the fact that in contrast to long lasting kisses or emotional speech, embraces take place frequently between platonic friends and even unfamiliar individuals (Forsell & Åström, 2012). Thus, the emotional condition might have lacked a strong affective component as embraces are not a partner-specific interaction eliciting strong emotional responses. Furthermore, embraces are usually shorter than the one-minute interval employed in our experiment (Nagy, 2011), which was necessary for reliable data acquisition. The unusually long duration could have negatively influenced our results in this experiment.

Given the relative novelty of the approach and the experimental paradigm, there are several limitations associated with the present study. On the one hand, the sample size of the study was rather low even for a within-subject design. Thus, it might have prevented the detection of smaller effects due to insufficient power. On the other hand, the present paradigm employed no negative emotional condition. Unfortunately, the lack of a negative emotional condition does not allow to conclusively embed our results into theories of emotional lateralization as the valence hypothesis distinguishes between positive and negative rather than positive and neutral affective states.

In conclusion, we found differences in alpha and beta asymmetries during emotional compared to neutral conditions in highly ecological situations that were in line with models of emotional lateralization integrating frontal valence and posterior right-hemispheric processing. To provide conclusive evidence in this regard, future studies should however conceive a similar experiment and include a negative emotional condition (e.g. listing annoying habits of the partner) that was not present in this study. Furthermore, the ecological validity of the present study is rather limited to couples from Western societies. For kissing, there are notable differences in both frequency and lateralization based on the context (Jankowiak, Volsche, & Garcia, 2015; Sedgewick & Elias, 2016; Sedgewick, Holtslander, & Elias, 2019), especially for cultural contexts (Karim et al., 2017). Similar results have been found for the cradling of children (Saling & Cooke, 1984). Thus, it would be interesting to see if cultural differences in social behaviour are reflected in altered neurophysiological processing of these types of interactions.

## Acknowledgements

This work was supported by the Deutsche Forschungsgemeinschaft grant number OC127/9-1 and the Research Training Group “Situated Cognition” (GRK 2185/1).

---

**SI Figure 1.**
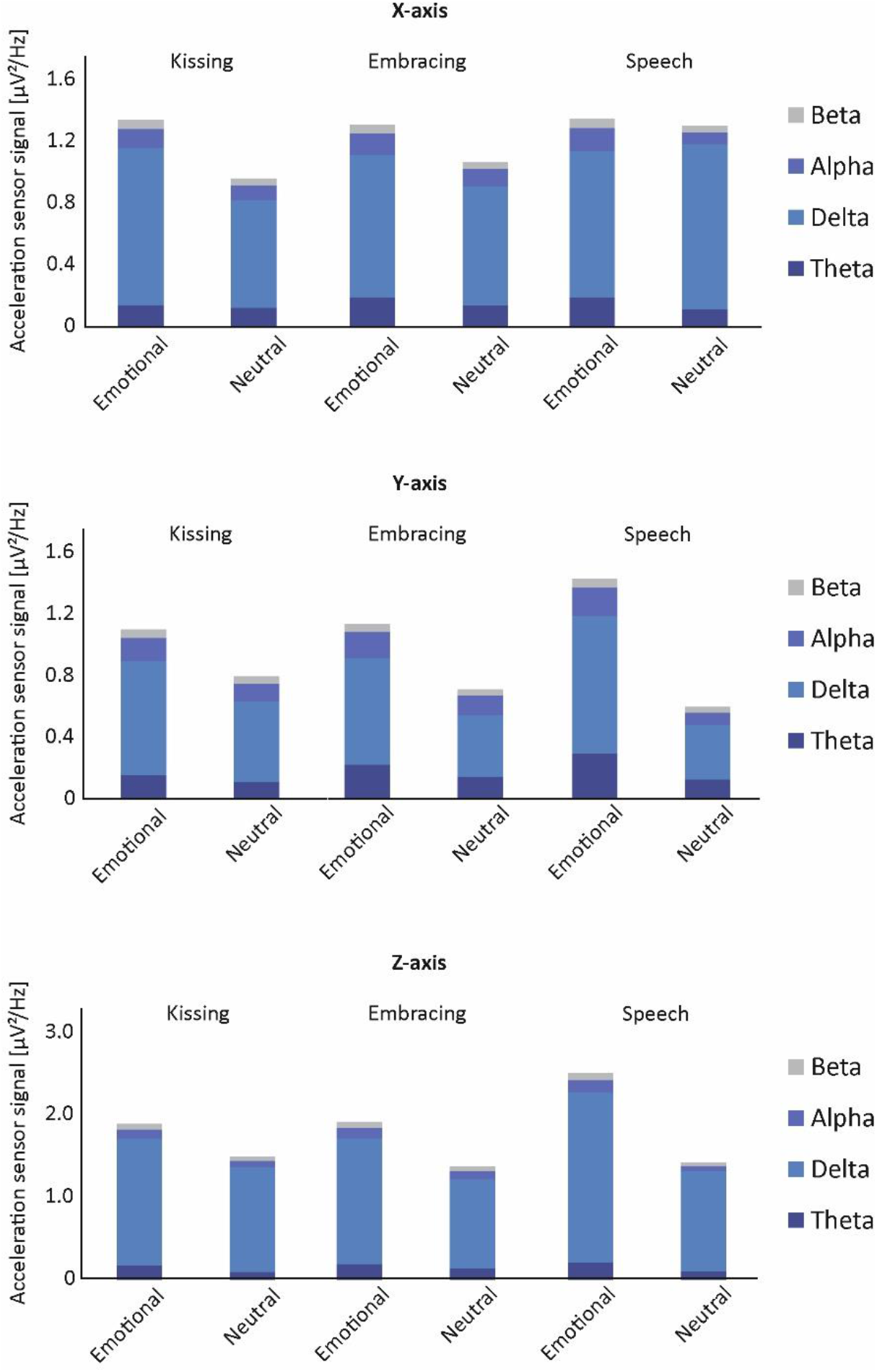
Head movements for the X- (top), Y- (center) and Z-axis acceleration sensors (bottom) across the three behavioral tasks and the relevant frequency bands. Depicted is the grand average across all participants for each individual behavior. Note that slow oscillations in the delta band dominated the activity measured by the acceleration sensors as head movements are generally slow.

